# Nanopore direct-RNA sequencing reveals TGEV epitranscriptomic and transcriptomic landscapes modulated by gene 7

**DOI:** 10.1101/2025.08.05.668584

**Authors:** Qingqiu Jiang, Zhihao Guo, Lu Tan, Yanwen Shao, Xiaomin Zhao, Haidong Wang, Runsheng Li

## Abstract

Viral non-structural proteins have gained increasing attention for their roles in regulating host-virus interplay and reported to act as a key mediator of host and virus RNA modification dynamics. Transmissible gastroenteritis virus (TGEV) gene 7 has been implicated in virulence, but its other molecular functions remain unclear.

Here, we compared wild-type TGEV (TGEV-wt) with a recombinant strain lacking gene 7 (TGEV-Δ7) in swine testis cells using Oxford Nanopore direct RNA sequencing. High-coverage datasets enabled simultaneous profiling of the full-length transcriptome, N6-methyladenosine (m6A) modifications, and polyA tail length.

Deletion of TGEV gene 7 halved viral RNA replication yet increased m6A modification by ∼32 % across the viral genome, and elevated host m6A levels by ∼17 %, accompanied by reciprocal shifts in the m6A regulators FTO (eraser) and RBM15 (writer). Despite bulk transcriptome changes were comparable between strains, gene 7 deletion introduced additional DEGs beyond WT infection, showing stronger enrichment of antiviral and chemokine pathways, indicating heightened innate immunity. PolyA analysis uncovered the polyA features of TGEV gRNA and sgRNAs, and revealed a gene 7 dependent extension of viral by 7 nt, but not host polyA tails. These findings highlight RNA-modification machinery as a potential target for coronavirus control and provide a framework for vaccine strategies exploiting gene 7 attenuation.

## INTRODUCTION

Transmissible gastroenteritis is an acute and highly contagious enteric disease affecting swine of all ages, characterized by vomiting, diarrhoea, and dehydration [1, 2]. In piglets, the mortality rate can be up to 100%, resulting in substantial economic losses in the swine industry [3, 4]. The causative agent is transmissible gastroenteritis virus (TGEV), an enveloped *Alphacoronavirus.* TGEV possesses a large, positive-sense single-stranded RNA genome of approximately 28.5 kb, comprising nine open reading frames (ORFs) arranged in the order of 5’ UTR-ORF1a-ORF1b-S-ORF3a-ORF3b-E-M-N-ORF7-3’ UTR [5, 6].

Following the entry of TGEV, the positive-sense genomic RNA (gRNA) is synthesized as a full-length negative template and serves as a template for the synthesis of progeny viral RNA [7]. Genomic RNA and subgenomic RNAs (sgRNAs) are used to translate structural proteins including spike (S), envelope (E), membrane (M), and nucleocapsid (N) proteins, which are essential for virion assembly and the viral infection process [8, 9]. In addition, TGEV encodes several non-structural and accessory proteins, including ORF1a, ORF1b, ORF3a, ORF3b, and ORF7 [10]. Notably, coronavirus non-structural proteins have been reported to influence viral virulence, host immune modulation, replication efficiency, and RNA modifications [11, 12].

N6-Methyladenosine (m6A) is the most abundant internal mRNA modification in eukaryotic cells, known to regulate mRNA stability, splicing, and translation efficiency [13, 14]. The abundance of host m6A is dynamically regulated by the writers (METTLs and RBM15), erasers (FTO), and readers [13]. Increasing evidence suggests that during infection, viruses induce alterations in host m6A, which in turn promote viral replication [15, 16]. And disruptions in host m6A writers pose a great impact on the survival of the virus [17, 18]. The m6A modification has become a key feature in studying virus-host interaction [19]. However, the detections of m6A sites in both host and viral RNAs are still hard for methods developed based on short-read sequencing [20]. Nanopore direct RNA sequencing records the current changes when the full-length RNA molecule passes through the nanopore, and the electrical signal can be converted into nucleotide sequence. In many cases, modified ribonucleotides will cause changes in ion signals and affect the accuracy of downstream base calls [21]. Several methods were developed exploiting changes in current or increased error rates to identify modified RNA sites [22]. The host DRACH m6A modification detection can reach 90% (m6Anet) [23]. And the viral RNA modification can also be profiled with a fine-tuned cutoff from a negative control [24].

The TGEV ORF 7 only contains one gene (gene 7) which encodes a non-structural protein consisting of 78 amino acids, with a molecular weight of approximately 9.1 kDa [10, 25]. Although not essential for virus viability, gene 7 has been reported to help the virus antagonize host antiviral responses [25, 26]. Also, gene 7 is directly linked to the 3’ untranslated region (UTR) of all TGEV sgRNA, suggesting a potential role in regulating their expression [27]. With the development of RNA modification detection technology, the RNA modifications in both the host and viral genomes could be profiled [15, 28]. However, the impact of TGEV gene 7 on RNA modification has not been systematically studied.

In this study, we employed Nanopore direct RNA sequencing to investigate the transcriptomic and epitranscriptomic landscapes of the host and virus, resulting from abrogation of TGEV gene 7. Our results provided a comprehensive overview of how TGEV gene 7 modulated m6A modification of both host and virus. Additionally, we found that the TGEV gene 7 significantly influences virus replication and the length of viral polyA tails. Furthermore, we identified related cellular pathways that were regulated by TGEV gene 7 through its impact on RNA modification systems. This result offers novel insights into the role of TGEV gene 7 in virus-host interactions. Generally, our findings underscore the importance of gene 7 in TGEV pathogenesis and suggest potential targets for antiviral strategies or vaccine design [29].

## MATERIALS AND METHODS

### Cells and viruses

Swine Testis cells (ST cells) were purchased from the American Type Culture Collection. ST cells were grown at 37°C with 5% CO_2_ in Minimum Essential Medium alpha supplemented with 10% fetal bovine serum, vitamins, L-glutamine, and penicillin-streptomycin.

We obtained wild-type TGEV (hereinafter referred to as TGEV-wt) and a recombinant TGEV which was deleted gene 7 (hereinafter referred to as TGEV-Δ7) from Northwest Agriculture and Forestry University.

### Virus infection and RNA isolation

ST Cells were seeded into 6-well plates at a density of 5 × 10^5^ cells/well and infected with viruses at MOI 1. After one hour of incubation, the cells were washed with phosphate-buffered saline three times and covered with a 1 mL cell culture medium, followed by further incubation. At 1-day post-infection (dpi), the supernatant was removed, and 800 μL TRIzol Reagent (Invitrogen, Waltham, MA, USA) was added to the cell monolayer. Phase separation and total RNA precipitation were performed according to the manufacturer’s instructions.

### Direct RNA sequencing (DRS) and data processing

Poly(A)+ RNAs were enriched from total RNAs using Dynabeads mRNA Purification Kit (Invitrogen). A total of 1000 ng of poly(A)+ RNAs were subjected to DRS library preparation using an SQK-RNA002 kit (Oxford Nanopore Technologies, Oxford, UK). The optional reverse transcription step was included using SuperScript III Reverse Transcriptase (Invitrogen). Sequencing was performed on the MinION platform using R9.4.1 flow cells (Oxford Nanopore Technologies).

Reads were basecalled using the Dorado workflow (v0.5.0) under the rna002_70bps_hac@v3 model (https://github.com/nanoporetech/dorado). The resulting FASTQ files were aligned to the TGEV genome (GenBank Accession Number: NC_038861.1) and *Sus scrofa* transcriptome (GenBank Accession Number: NC_010443.5) using Minimap2 v2.17 with parameter settings “-ax map-ont” [30]. Mapping results were stored in SAM files, which were subsequently converted into bam files and sorted and indexed using SAMtools v1.17[31]. Qualification control information was extracted using a custom Python script based on the result from SeqKit v2.9.0 and Giraffe v0.2.3[32, 33]. Reads mapped to the host in the sample of healthy ST cells without infection will be referred to as ST-mock, in the sample of ST cells infected with TGEV-wt will be referred to as ST-wt and in the samples infected with TGEV-Δ7 will be referred to as ST-Δ7. Reads mapped to TGEV will be referred to as TGEV-wt and TGEV-Δ7 according to the type of virus infected.

### Current feature showcase with nanoCEM

nanoCEM (version 0.0.6.3) was utilized for displaying Nanopore sequencing current at site level[34]. All raw data were transformed into blow5 format by blue-crab (https://github.com/Psy-Fer/blue-crab)[35]. We selected the f5c_ev mode with the options “--norm”, “--rna” and “--pore r9”.

### Detections of m6A sites

m6Anet (v1.1.1) relies on the eventalign module in Nanopolish (v0.14.0) to assign raw current signals to each nucleotide[23]. The signal features, including mean, standard deviation, and signal length, are used as input to a Multiple Instance Learning-based neural network model. We incorporated mod_ratio values computed as m6A level for each DRACH in our analysis.

For the comparative methods, we applied a python script to merge the results from Tombo and xPore (https://github.com/lrslab/Scripts_merge_DRS_methods)[20].

1. Tombo_com v1.5.1 is the canonical sample comparison method provided by Tombo. It offers two ways for modified base detection, “model_sample_compare” and “level_sample_compare”. The latter approach was adopted in the present study. The “--store-p-value” option was applied to save P-values for subsequent analysis. The “text_output” option was employed to extract P-value and Difference statistics.
2. xPore v2.1 employs the outputs derived from the Nanopolish eventalign module for all samples to generate a configuration file. The prediction of modified positions was determined by P-value and differential modification rate from the statistical test results.

#### Cutoff selection

Reads of ST-mock were divided into two subsamples randomly, then employed a two-tailed 99% confidence interval to establish comparative cutoffs (plus or minus 0.330 in this study). This cutoff was used to identify the most significantly changed m6A sites

### DEG and DEI comparison analysis

A gene was classified as a differentially expressed gene (DEG) between tissues when meeting two criteria: (1) transcript abundance (measured in reads per million) demonstrated over two-fold change (|log_2_FoldChange| > 1), (2) the False Discovery Rate (FDR) <0.05. Statistical analysis was performed using the edgeR package (version 3.30) in R, with FDR calculations conducted through Benjamini-Hochberg correction following quantile normalization of read counts[36]. A single gene can form different isoforms during transcription through alternative splicing, which is an essential post-transcriptional mechanisms[37]. A transcript isoform was classified as differentially expressed (DEI) between two tissues if the alteration in its relative abundance (calculated as the percentage of its read count relative to all transcripts within the same genetic locus) exceeded 20%[38]. We use TrackCluster (version 0.1.7) to identify isoforms [39]. Then, ClusterProfiler package was used to deal with Gene Ontology Enrichment Analysis to explore the enrichment of certain functions or features in the DEG and DEI gene sets, respectively[40]

### Viral gRNA and sgRNAs expression

After virus entry into cells, genomic RNA (gRNA) will serve as the initial template to combine a nested set of subgenomic RNAs (sgRNAs) with a same 5’ UTR leading[41]. SgRNAs always start from a certain ORF of the virus and completely contains the remaining sequence of the viral genomic RNA until the 3’ polyA tail. But only the ORF closest to 5’ UTR would be translated, whether genomic RNA or subgenomic RNAs[41]. Therefore, we divided the mRNA mapped to the viral genome into gRNA and different sgRNAs according to the viral ORF to which the mRNA 5’ belongs. Considering that nanopore sequencing starts from the 3’ end, the 5’ end sequence may be degraded during the process, resulting in incomplete ORF. After deleting the polyA tail, we only require counting from the 3’ end of the reads, and at least one base after that belongs to the corresponding ORF. The production of ORF1b depends on the ribosomal frameshift that occurs after the production of ORF1a, so ORF1b does not generate new sgRNAs[42].

Instead, it forms a larger protein together with ORF1a, so it is still expressed using gRNA as when ORF1a is expressed alone. In addition, it should be emphasized that the mutation at the front end of TGEV gene 7 will only cause the protein encoded by gene7 to fail to be translated and will not affect the production of sgRNA of gene 7. After CPM, we used the relative expression of ORFs to compare the effect of gene7 abrogation on the expression of viral ORFs.

### Poly(A) length estimate

Nanopolish (version 10.2) was used to calculate poly(A) length of each read with command “nanopolish polya” (https://github.com/jts/nanopolish)[43]. The length of the polyA was inferred from the dwell time of the raw ion signal at the unaligned part of the 3’ end of the reads.

We only retained reads with polyA length ≤ 250 nt for further analysis. We first performed kernel density estimation on the data of gRNA and each sgRNA to identify peaks in the distribution of polyA tail length. The local maximum (peak) of the density curve was detected as a point with higher density than the adjacent points. This process was implemented in R using the density() function (adjust = 1.2). To reduce false positives caused by small fluctuations, only peaks with a signal-to-noise ratio greater than 1 were considered.

## RESULT

### Direct RNA Sequencing obtained enough reads for ST cell and TGEV transcriptome analysis

ST cells were infected at MOI 1 with TGEV or TGEV-Δ7. At 17h post-infection, when the virus reached its highest replication titer, polyadenylated RNAs were isolated from infected cells and sequenced via ONT direct RNA sequencing. The healthy ST cell without infection was also collected and sequenced as mock (Figure 1A).

**Figure 1.**
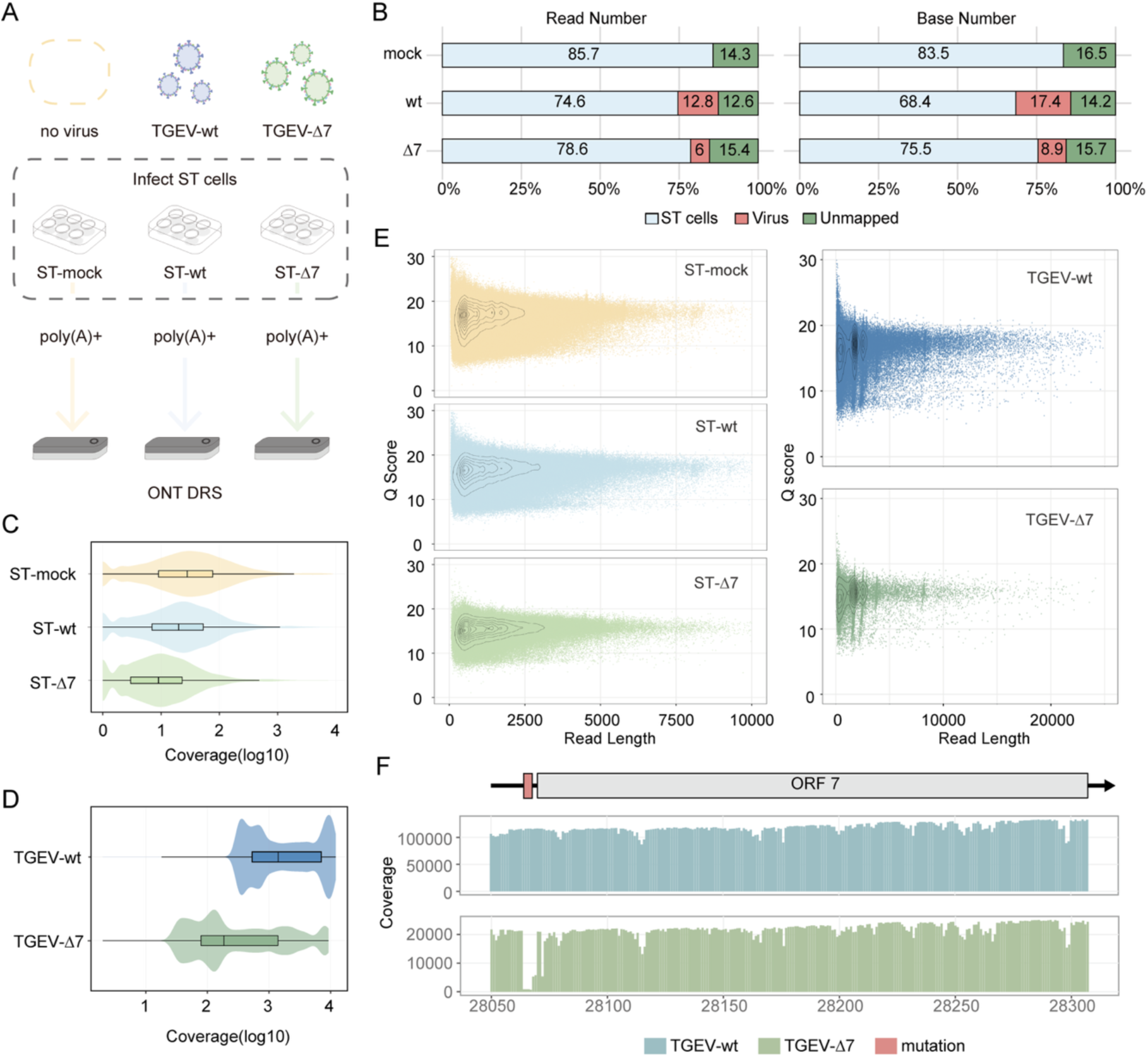
Raw read characterization and mapped reads analysis. (A) Comparative experimental design and sequencing process of each group. (B) The proportion of host reads and viral reads from the three sequenced samples. (C) Gene coverage of reads mapped to the swine genome. (D) Genes coverage of reads mapped to the viral genome. (E) Distribution of Q scores for different read lengths. As the read length increases, the Q scores of both viruses remain at a high level, with no significant downward trend. (F) Coverage bar plot around TGEV gene 7 showed a frameshift mutation caused by a three-base deletion at the front end.

After basecalling, we obtained 1,918,474, 1,248,609 and 486,170 reads from mock, TGEV-wt infected sample, and TGEV-Δ7 infected sample, respectively. Over 85% of reads from each sample could be mapped to the swine genome or TGEV genome, with no TGEV mapped reads in mock, confirming the absence of pre-existing virus contamination (Figure 1B, S1). What’s more, viral RNA abundance differed markedly between infections, with the proportion of TGEV-wt being two times higher than TGEV-Δ7 (12.8% vs 6% in read number or 17.4% vs 8.9% in base number, Figure 1B), confirming TGEV gene 7 was a key virulence factor. Host-derived mRNA reads were hereafter designed as ST-mock (uninfected), ST-wt (TGEV-wt infected), and ST-Δ7 (TGEV-Δ7 infected), while viral reads were labeled according to their respective strains (TGEV-wt or TGEV-Δ7).

Host transcriptome sequencing achieved comprehensive genomic representation, with over 95% of annotated swine genes detected across samples (Figure 1C). Viral sites showed robust coverage, maintaining a minimum median depth of 50×, which is enough for reliable direct RNA sequencing-based RNA modification analysis (Figure 1D) [23]. All groups demonstrated consistent sequencing quality, with median Q-score of 16.65 (ST-mock), 16.47 (ST-wt), and 15.28 (ST-Δ7) for host reads, and 16.51 (TGEV-wt) and 15.26 (TGEV-Δ7) for viral reads. In addition, the GC content of TGEV-wt in the viral sequencing results was 38.24%, and the GC content of TGEV-Δ7 was 38.08%, which is consistent with the overall low GC content characteristics of the Coronaviridae family [44] (Table 1). Viral read length distributions revealed four peaks at 1500 nt, 2500 nt, 3800 nt and 8000 nt, which originated from expressions of sgRNAs encoding nucleoprotein, envelope protein, non-structural protein 3 and spike protein (Figure 1E, S1, Table S1). We checked coverage per site and found that nanopore DRS can capture the three-base mutation in front of gene 7 as described in the previous article, determining the correct abrogation of TGEV gene 7 in TGEV-Δ7 sample (Figure 1F).

**Table 1.** Raw data feature based on Dorado basecalling.

|  | Number of reads | Average length | N50 | Median Q score | GC(%) |
| --- | --- | --- | --- | --- | --- |
| ST-mock | 1,643,837 | 1,194.7 | 1,609 | 16.65 | 50.78 |
| ST-wt | 931,761 | 1,412.1 | 1,923 | 16.47 | 49.15 |
| ST-Δ7 | 382,124 | 1,409.9 | 1,861 | 15.28 | 49.5 |
| TGEV-wt | 159,496 | 1,991.6 | 2,574 | 16.51 | 38.24 |
| TGEV-Δ7 | 29,227 | 2,041.2 | 2,544 | 15.26 | 38.08 |

Our nanopore direct RNA sequencing approach generated high-coverage, quality-controlled datasets across all experimental groups, and confirmed the loss of function mutation of gene7 in TGEV-Δ7 virus while ensuring comparability between wild-type and mutant datasets. This foundational data quality enabled system-level analysis of transcriptomic and epitranscriptomic alterations and mechanistic dissection of TGEV gene 7’s function in modulating RNA processing.

### Inverse regulation of host m6A methylation landscapes between TGEV-wt and TGEV-Δ7 infection

Around 230,000 DRACH sites were detected by m6Anet in swine transcriptomes (Figure 2A). We performed pair-wise comparisons of ST cell m6A proportion and selected the shared m6A sites that can be profiled (DRACH motifs with read coverage >= 5) in each pair for downstream analysis. To identify the most significantly changed m6A sites, we randomly divided reads of ST-mock into two subsamples and applied a two-tailed 99% confidence interval to determine the significance cutoffs as 0.330.

**Figure 2.**
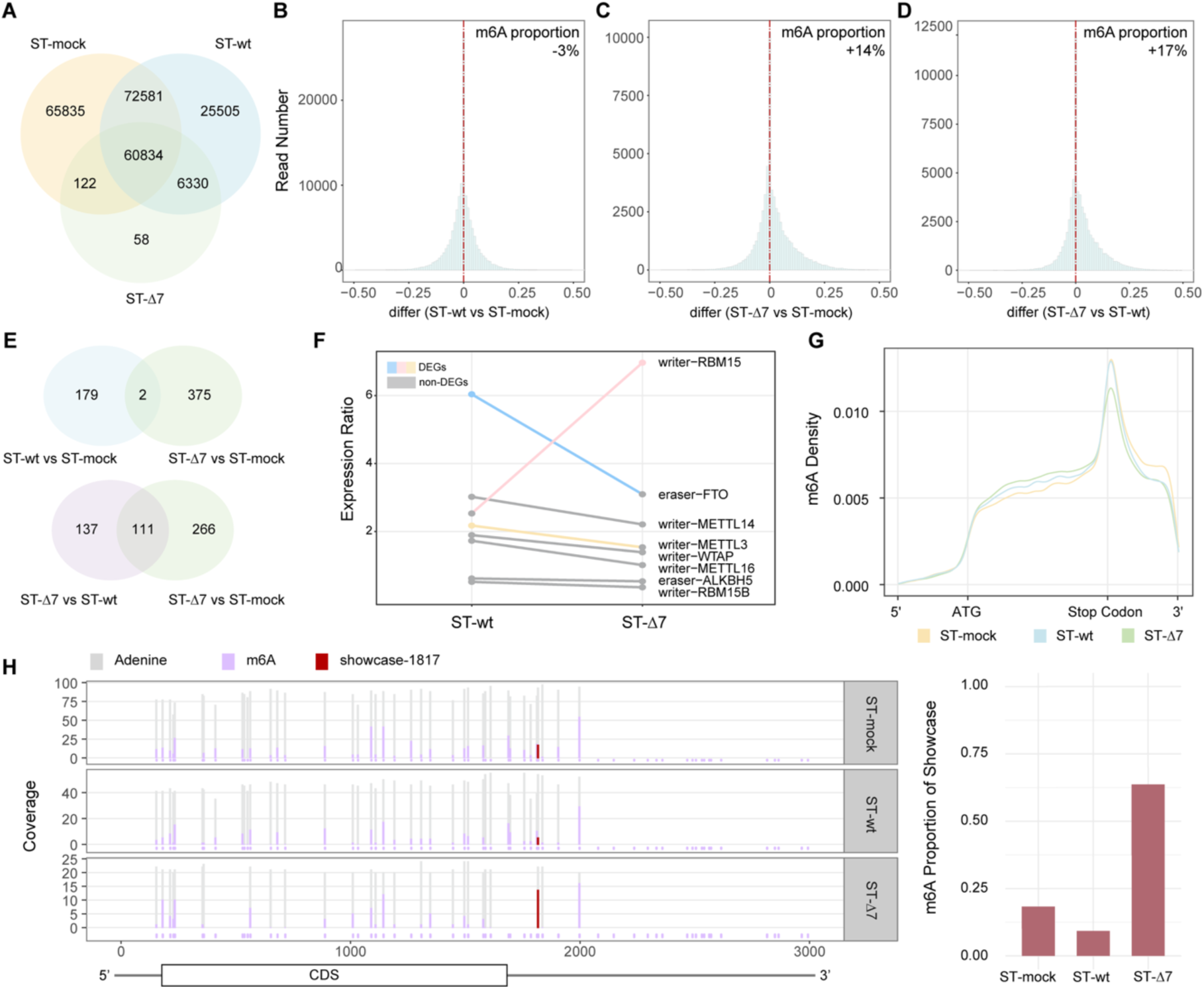
Overall analysis of m6A modification in ST cells. (A) m6A sites (DRACH sequence) of host (ST cell) detected by m6Anet. The overall difference of m6A levels between ST-wt and ST-mock (B), ST-Δ7 and ST-mock (C) and ST-Δ7 and ST-wt (D). (E) The overlapping of significantly changed m6A sites between pairwise comparisons. Here we require not only the same site, but also the same modification changes at the site. (F)Expression ratio of genes regulating m6A levels compared with ST-mock. Colored lines indicate that the gene is a statistically differentially expressed gene in at least one group, and the points of that group are also colored, otherwise they are gray. (G) m6A site density in ST cell mRNA. (H) Methylation proportion of each m6A site around gene SNX1. The dark red site (located at base 1817 of gene) is used as a showcase to show the changes in the proportion of m6A in different samples.

During TGEV-wt infection, the overall proportion of m6A modifications in mRNA decreased (Figure 2B). Compared to ST-mock (healthy, uninfected ST cells), 52,567 m6A sites in ST-wt were increased, while 62,495 sites showed reduced m6A levels. On average, the global m6A level decreased by approximately 3%. By applying a threshold of >0.330 to identify significantly changed sites, we found that ST-wt sample had 67 sites with significantly higher m6A modification, and 114 sites with significantly lower modification (Table S2).

The ST cell infected with TGEV-Δ7 showed a different trend of overall m6A modification change. When compared with ST-mock, we found that 31,303 m6A sites in ST-Δ7 showed higher m6A modification proportion, while 23,204 sites were lower (Figure 2C). The overall increase of m6A proportion after TGEV-Δ7 infection was 14%. For the selection of significantly changed sites, 332 sites were significantly higher and only 42 sites were significantly lower (Table S3).

When comparing the differentially infected ST cells, the difference in m6A proportion became more obvious. There were 36,786 m6A sites in ST-Δ7 that had higher m6A proportion, while 22,238 sites in ST-wt had higher m6A proportion (Figure 2D). The overall increase of ST cells m6A modification between TGEV-Δ7 infection and TGEV-wt infection was 17%. For the significantly changed m6A sites, we detected 239 sites that had significantly higher modification, and 9 sites had significantly lower modification in ST-Δ7 when compared to ST-wt (Table S4).

Analysis of significantly changed sites also showed that when using ST-mock as the background, the m6A modification of the two infection groups was very different. There were only two overlaps of sites with the same change trend in the two infected samples, and the similarity of changes caused by different infections was very low. Compared with ST-wt or ST-mock, 111 points of ST-Δ7 changed in the same direction, indicating that deletion of TGEV gene 7 dominated the production of specific epigenetic responses (Figure 2E). TGEV gene 7 had a strong regulatory effect on host m6A modification over infection.

Refer to writer genes responsible for m6A methylation and eraser genes responsible for m6A demethylation that had been identified [13]. We examined the expression fold of writers and erasers in ST-wt and ST-Δ7 by using gene expression of ST-mock as the baseline. m6A modifications regulators in ST-wt and ST-7 were activated to various degrees. Fat mass and obesity associated gene (FTO, eraser) was significantly upregulated in ST-wt by two times higher than ST-Δ7. In contrast, in ST-Δ7, Putative RNA-binding protein 15 (RBM 15, writer) was significantly upregulated by 2.6 times higher than ST-wt (Figure 2F). The expression fold of the regulatory gene explains the observed decrease in methylation levels in ST-wt and increase in ST-Δ7m6A.

Even though the m6A proportion changed greatly, the distribution of modification sites on CDS remained stable. All ST cells showed enrichment of m6A modification near the stop codon (Figure 2G), which is consistent with the distribution of m6A modification in mammals [45]. Most of the measurable points near the stop codon were consistent with the trend that ST-Δ7 was greater than ST-mock and ST-wt. For example, the m6A site close to the stop codon of the gene SNX1 showed a dramatic increase in ST-Δ7 (Figure 2H).

### TGEV-Δ7 exhibited enhanced viral RNA m6A methylation compared to TGEV-wt

Viral RNA modification differences between TGEV-wt and TGEV-Δ7 were observed using m6Anet (Figure 3A-C) and a pipeline integrating Tombo and xPore as a secondary verification method (Figure S3A-C). Comprehensive viral m6A profiling revealed systematic hyper modification in TGEV-Δ7.

**Figure 3.**
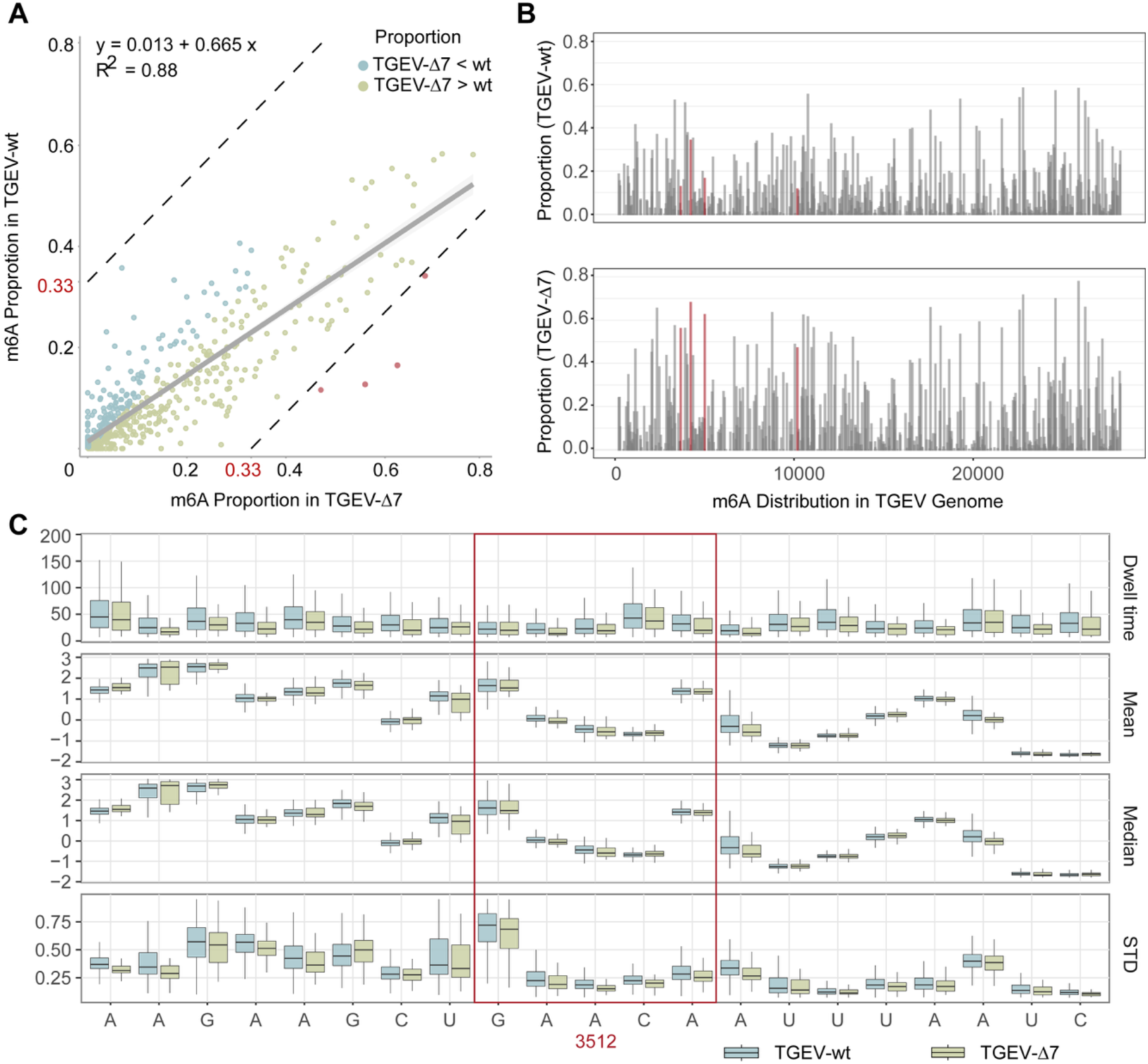
Comparison of the m6Asite in viral RNA detected by m6Anet and the intersection of xPore and Tombo. (A) RNA m6A modification proportions detected by m6Anet. The correlation of the m6A proportions between TGEV-wt and TGEV-7 was examined by linear regression. The significantly differed m6A sites are colored in red. (B) Genomic distribution of m6Anet detected modification sites in the viral genome. The four significantly differed m6A sites are highlighted in red. (C) The ONT raw current changes of the m6A site at position 3512. The red frame highlighted was DRACH site (GAACA).

In total, 588 m6A sites with DRACH motif could be profiled using m6Anet. Among them, 336 sites showed higher modification proportion in TGEV-Δ7, when compared with TGEV-wt. Only 201 sites displayed a reversed trend (Figure 3A). The summed proportion from all m6A sites in TGEV-Δ7 was 32 % higher than that in TGEV-wt (Figure 3A and B).

Although there was no evidence of motif spatial clustering in viral genome. We identified the four most significantly differed sites with a threshold of m6A proportion difference above 0.330 (Figure 3A). All four sites were located in ORF1a (Figure 3B) and exhibited elevated m6A in TGEV-Δ7 (Figure 3A and B). The m6A modification at the DRACH motif would likely lead to a decrease in raw current signal than the canonical A [46]. When using NanoCEM to compare the raw current signals differences around one significantly differed site. We showed the current around site 3512 here, the decreased current levels around site 3512 could be observed to validate the change of m6A proportion (Figure 3C). The results of xPore and Tombo were combined to obtain 344 sites with potential modification differences (Figure S3). These 344 sites were further combined with the results of m6Anet, and only 23 common sites were found. According to the results of m6Anet, 16 sites showed a higher proportion of m6A modification in TGEV-Δ7 than in TGEV-wt (Table S4).

TGEV-wt and TGEV-Δ7 also showed a similar m6A change trend as the host. In the comparison of the infected groups, we found that the m6A in both host and viruses increased in the infected group lacking TGEV gene 7. Suggesting that TGEV employed gene 7 to avoid the host m6A modification, as a deletion of gene 7 would reverse the effect.

### Conserved transcriptomic profiles with enhanced host antiviral resistance in TGEV-Δ7 infection

We used TrackCluster to quantify the host transcriptome. Stringent thresholds (genes: total counts >15; isoforms: total counts > 6) were applied to confirm the expression of genes and isoforms. Correlation analysis revealed high similarity between ST-wt and ST-Δ7 (Figure 4A).

**Figure 4.**
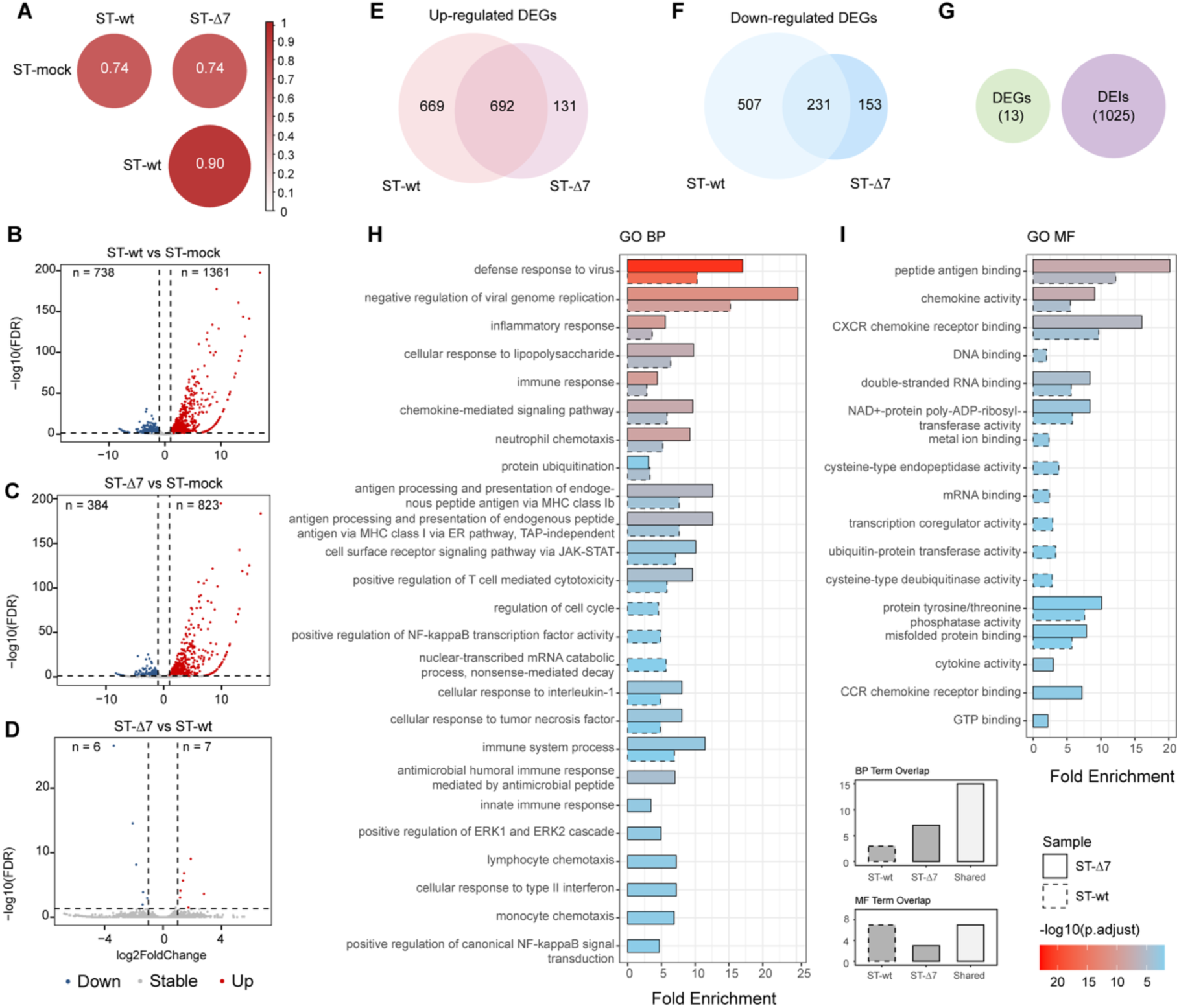
Comparative analysis of host differentially expressed genes (DEGs) and differentially expressed isoforms (DEIs). (A) Spearman correlation analysis of transcriptome response between healthy ST cells (ST-mock), ST cells infected with TGEV-wt (ST-wt), and ST cells infected with TGEV-Δ7 (ST-Δ7). Volcano plots of DEGs (|log_2_FC| ≥ 1, FDR < 0.05) gained comparison between ST-wt and ST-mock (B), ST-Δ7 and ST-mock (C), ST-Δ7 and ST-wt (D). The DEGs obtained by comparing the two different infection groups with the mock group were overlapped, and both up-regulated DEGs (E) and down-regulated DEGs (F) showed significant intersection. (G) Overlap of DEGs and DEIs between ST-wt and ST-Δ7. Gene Ontology (GO) enrichment analysis of up-regulated DEGs with p-adjust < 0.01, Biological Process (H) and Molecular Function (I) were displayed separately. Cellular Component have no enrichment in any term.

Differentially expressed gene (DEG) analysis showed TGEV-wt infection induced more pronounced changes. When compared with ST-mock, ST-wt showed a total of 1361 significant up-regulated genes, and 738 significant down-regulated genes (Figure 4B). While ST-Δ7 showed only 823 up-regulated genes and 384 down-regulated genes (Figure 4C). Direct comparison between ST-Δ7 and ST-wt showed minimal difference with only 13 DEGs (Figure 4D). Although ST-wt versus ST-mock showed more DEGs compared with ST-Δ7 versus ST-mock, more than half of the DEGs obtained in the two comparisons were the same. (Figure 4E, F). Notably, while gene-level changes were limited, we observed numerous differentially expressed isoforms (DEIs) in ST-wt versus ST-Δ7, indicating isoform-level regulation could be independent from gene level (Figure 4G).

We then used DEGs for Gene Ontology Enrichment (GO) analysis and found that upregulated DEGs in ST-wt and ST-Δ7 showed similar enrichment patterns in the “biological process” and “molecular function” categories, while downregulated DEGs showed similar enrichment patterns in the “cellular component” category (Figure 4H-J).

Compared with ST-mock, most of DEGs enriched in biological process were up-regulated and exhibited striking convergence where ST-wt was enriched in 18 pathways and ST-Δ7 was enriched in 22 pathways, with 15 pathways were shared. Both ST-wt and ST-Δ7 showed strong antiviral immunity and inflammation themes, but the enrichment fold and significance of ST-Δ7 were generally higher (for example, ST-Δ7’s “defense response to virus” Fold Enrichment=17.05 vs. ST-wt’s10.29), and ST-Δ7 contained more chemotaxis-related terms (such as “neutrophil chemotaxis”) (Figure 4H). This suggests that the immune response of ST-Δ7 is more intense and may involve a wider range of cell types (such as Lymphocyte chemotaxis). The only enriched term among the downregulated DEGs involved mitochondrial cytochrome c oxidase assembly, indicating that mitochondrial energy metabolism or oxidative phosphorylation may be inhibited.

In molecular function (MF), the significantly enriched terms were all concentrated in up-regulated DEGs and related to viral defense and inflammatory signals. In general, ST-wt versus ST-mock was enriched in 14 pathways and ST-Δ7 versus ST-mock was enriched in 10 pathways, of which 7 pathways were shared by the two groups. Both samples were enriched for antigen presentation (SLA family) and chemokine-mediated immune cell recruitment, indicating similar basic immune activation mechanisms. ST-Δ7 had enhanced antiviral recognition: the enrichment fold of dsRNA binding function increased by 50%, and the OASL gene was only expressed in ST-Δ7 but not ST-wt. Inflammatory signals were also expanded, with new cytokine activity (IFN/IL6/TNF) and CCR chemotaxis. For ST-Δ7, the p-values of the same functions were lower and the enrichment folds were higher.

Analysis of MF also indicated that TGEV-Δ7 could lead a more active immune response, involving a wider range of inflammatory mediators (cytokines) and more efficient viral RNA recognition mechanisms (dsRNA binding).

Overall, gene enrichment analysis showed that TGEV-Δ7 infection triggered a stronger host immune response than TGEV-wt infection. This suggests that gene 7 of TGEV has the function of suppressing antiviral immunity. Consistent with this, TGEV-Δ7-infected cells consistently showed significantly higher enrichment of upregulated DEGs in antiviral and inflammatory pathways compared with TGEV-wt infection.

### Sharp decline in viral overall RNA replication with stable relative gene expression

The number of viral reads was normalized by CPM and aligned with the annotated viral genome to obtain the distribution map of virus reads on the genome. The step-like distribution is consistent with the characteristics of viruses using genomic RNA (gRNA) and subgenomic RNAs (sgRNAs) for transcription (Figure 5A, S2). We noticed that while the overall trend was similar, the expression level of TGEV-Δ7 could only reach half of that of TGEV-wt (Figure 5A). Viral mRNAs could be grouped based on the characteristics of TGEV gRNA and sgRNAs (Figure 5B). As gRNA and sgRNAs only translate the gene closest to the 5’ UTR during translation, we grouped reads according to which ORF the 5’ end base belongs to after mapping to the viral genome (Figure 5B). We found that the relative fold change of gene expression between TGEV-Δ7 and TGEV-wt was all close to 1, no difference could be observed (Figure 5C). The abrogation of TGEV gene 7 would not alter the expression ratio of viral genes.

**Figure 5.**
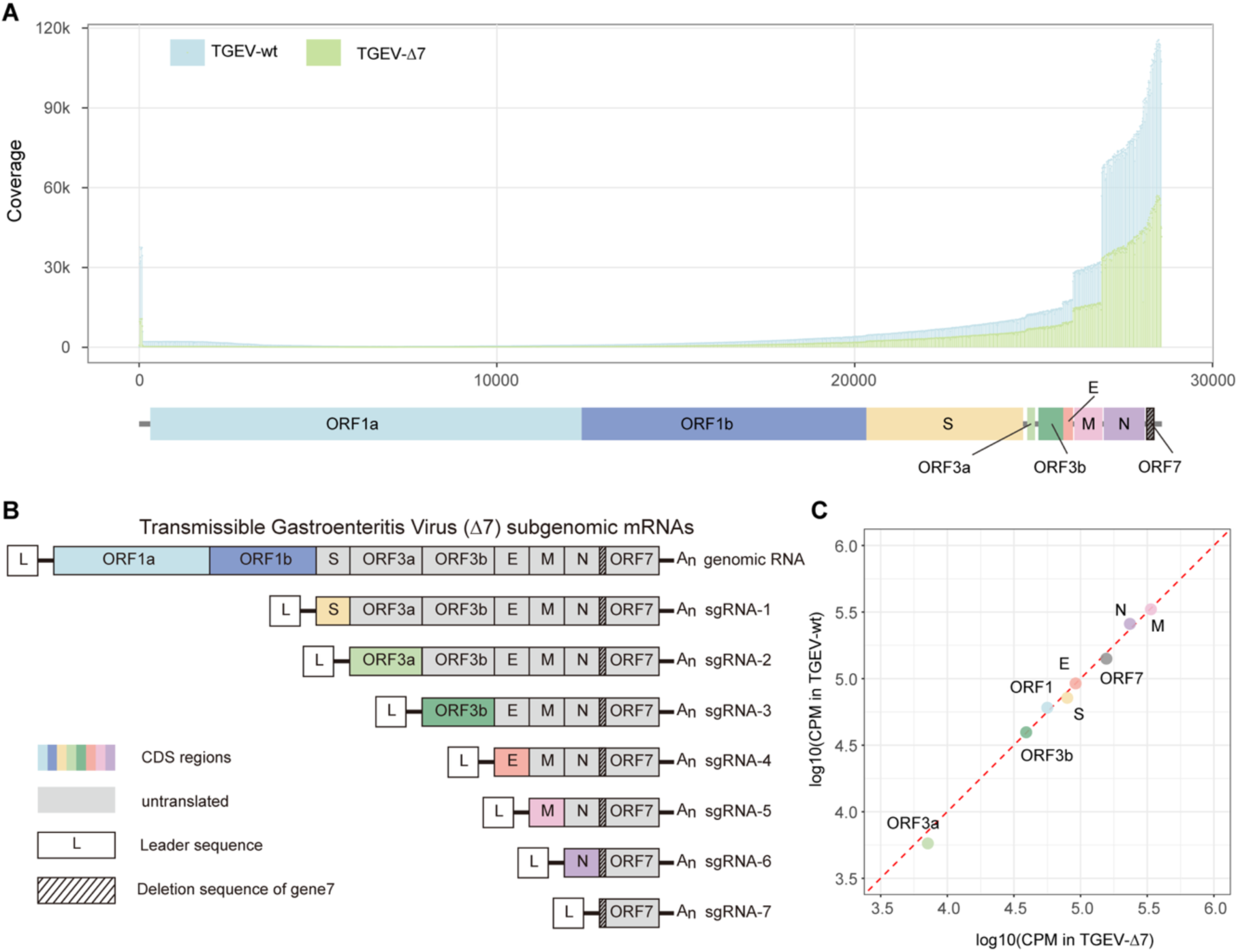
Relative expression of viral genomic RNA and subgenomic RNAs. (A) Coverage of each site of sequencing data aligned to TGEV reference. Blue bars represent TGEV-wt (above), and green bars represent TGEV-Δ7 (below). The X-axis is site location in the virus genome, and the y-axis is site coverage after normalizing (coverage/all reads * 10^6^). (B)Genomic and subgenomic RNAs produced during TGEV infection. Three bases were deleted at the beginning of the sequence of ORF7, indicated by the dotted area, which prevents gene7 from being successfully translated (C) Correlation of genomic and subgenomic RNAs expression between wild type TGEV (TGEV-wt) and recombinant TGEV which abrogated gene 7 (TGEV-Δ7), each point represents the log10(CPM) value of one gRNA or sgRNA.

### Deletion of gene 7 leads to a significant extension of the viral polyA tail during infection

We used genes with average CPM >= 5 to carry out the polyA tail analysis. Firstly, all reads of genes that pass the CPM filter were used for analysis. The median polyA length of ST-mock was ∼105. ST-wt showed a significant extension of ∼3nt, while the change in ST-Δ7 polyA length was not statistically significant (Figure 6A). TGEV-wt infection would prolong the mean length of polyA tail, but abrogation of TGEV gene 7 alleviated this situation.

**Figure 6.**
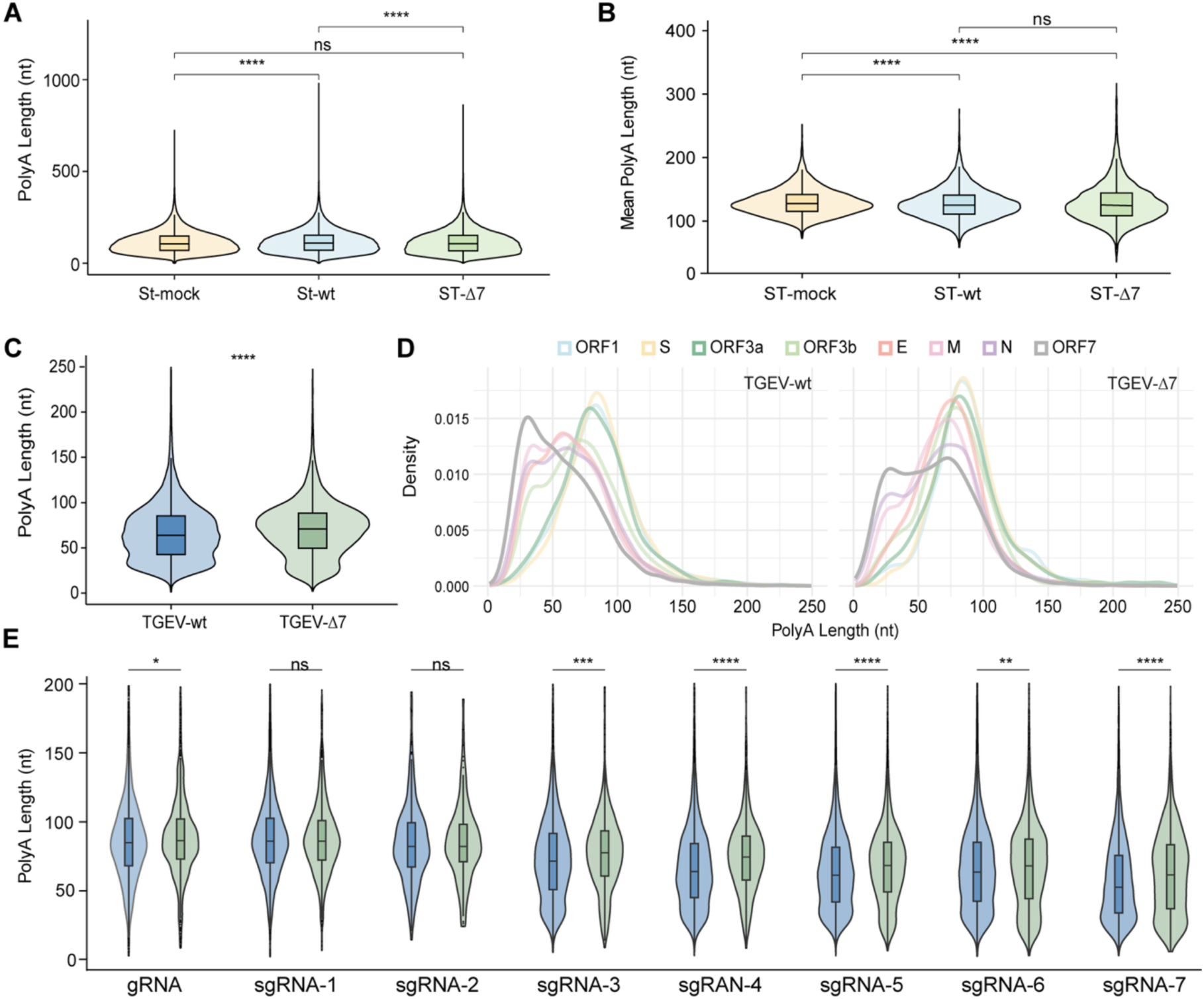
Changes in RNA polyA lengths of viruses. (A) PolyA length distribution of total mRNA from ST-mock, ST-wt, and ST-Δ7. (B) Mean polyA length distribution of filtered genes. (C) PolyA length distribution of all viral RNAs. (D) PolyA length distribution of viral reads after grouping according to genomic and subgenomic transcriptional features. (E) Pairwise comparison of TGEV-wt and TGEV-Δ7 genomic RNA and sgRNAs lengths. The Mann–Whitney U test was used to find significance

Recent studies have shown that short-tailed transcripts tend to be highly expressed [47]. To examine if the gene expression had an impact on polyA length, we separated the reads from each gene to calculate gene level polyA length. We found that the median polyA length of ST-mock was ∼128nt, and the median polyA lengths of ST-wt and ST-Δ7 were both ∼125nt (Figure 6B, Table S5). The polyA tail distribution results of ST-wt and ST-Δ7 are basically the same (Figure S4). Generally, the polyA tail length of the host cell showed a slight decrease after infection, but TGEV gene 7 did not bring additional effects.

We also counted the polyA length from viral transcripts. Measured against SARS-CoV2, TGEV-wt had a relatively long polyA tail, with a median length of 64 nt [48]. And the abrogation of TGEV gene 7 induced a significant polyA tail increase of 7 nt (Figure 6C). For gRNA and sgRNAs, in TGEV-wt, there were two polyA tail populations in sgRNAs used to translate ORF3b, M, N. The minor peak was around 33 ∼ 37 nt, while the major peak was around 60 ∼70 nt. However, the polyA length distribution for sgRNA-7 (translate ORF7) were different from all other sgRNAs, with the highest peak at only 30 nt (Figure 6D).

When compared with TGEV-wt, TGEV-Δ7 exhibited an additional major peak in sgRNA-7 at 73 nt, while the minor peaks of sgRNA-3 and −5 disappeared (Figure 6D, Table S6). Compared with TGEV-wt, most polyA tails of TGEV-Δ7 gRNA and sgRNAs increased by 3 ∼ 10 nt except for sgRNA-1 and sgRNA-2 (Figure 6E and Table S7). Notably, the overall polyA length seemed to align with the length of gRNA and sgRNAs (Figure 6E, spearman correlation R of TGEV-wt = 0.95, TGEV-Δ7 = 1). This trend leaded to a huge difference in the polyA length between gRNA and sgRNAs (Table S7). For example, the difference between TGEV-wt gRNA and sgRNA-7 reached 32.5 nt.

The different trend of host and viral polyA tail between TGEV-wt and TGEV-Δ7 infected samples indicated that TGEV had a polyA tail regulation mechanism independent from the host. The coronavirus protein translation efficiency was positively related to polyA length [49]. The overall polyA tail extension in TGEV-Δ7 would likely alleviate the adverse effects of the sharp decrease in TGEV-Δ7 RNA replication by improving the efficiency of translation.

## DISCUSSION

By using nanopore direct RNA sequencing, we demonstrated a link between TGEV nonstructural protein gene 7 and m6A modifications. Infection with TGEV-wt significantly upregulated the m6A eraser FTO, resulting in a 3% reduction in globe host m6A levels. While infection with TGEV-Δ7 robustly induced the m6A writer RBM15, elevating host m6A by a 14%. GO analysis of DEGs revealed that loss of TGEV gene 7 enhanced antiviral and chemokine pathways, and the resultant attenuation was further reflected by a shape decline in viral RNA replication. Finally, gene 7 deletion produced virus-specific extensions of polyA tails that contrasted with the host polyA length during infection.

While this high-throughput study offered innovative insights, we acknowledged several limitations. First, the absence of biological replicates and the comparatively low sequencing depth for TGEV-Δ7-infected ST cells could have introduced biased. Second, without in vitro–transcribed, unmodified RNA as a negative control, we were unable to validate the accuracy of the results of m6Anet-identified viral m6A sites. Finally, our xPore-Tombo pipeline couldn’t quantify absolute changes in modification stoichiometry, and its sensitivity remained uncertain without a corresponding unmodified reference.

We proposed that the diminished replication observed in TGEV-Δ7 was driven by the upregulation of m6A levels. Supporting this notion, suppressing host FTO, the m6A eraser, has been shown to impair replications of several RNA viruses [15]. Accordingly, the loss of gene 7 likely reduced viral replication indirectly by tilting the host epitranscriptomic balance toward increased m6A. The accumulation of m6A modification could also modulate innate immunity, as documented in SARS-CoV2 infection [48]. Our data indicated that gene 7 acted as an immune-evasion factor by simultaneously down-regulating the writer RBM15 and up-regulating the eraser FTO, thereby preventing excessive m6A on both host and viral RNAs and safeguarding viral replication. This mechanism may explain why TGEV infection lowered host m6A, whereas beta coronavirus infections typically elevated it [7].

Contrary to the negative correlation between polyA length and RNA modification reported previously [48]. Our study showed a direct positive correlation between polyA tail extension and increased m6A levels in TGEV-Δ7. This discrepancy may stem from fundamental differences in the predominant RNA-modification motifs examined. The SARS-CoV-2 study focused on the AAGAA motif [48], whereas our analysis targeted the canonical DRACH consensus for identifying m6A sites. Taken together, these observations broaden the framework for antiviral strategies that target RNA stability or epitranscriptomic pathways. Future research should dissect motif specific links between m6A and polyA regulation and evaluate the universality of these mechanisms across viral taxa.

## DATA AVAILABILITY

The raw nanopore sequencing data (FAST5 files) generated in this study have been deposited in the BioProject under the primary accession code PRJNA1256614.

## ACKNOWLEDGMENTS

This research was funded by the Early Career Scheme (CityU 21100521) and General Research Fund (Project No. 11105524) of the Research Grants Council of the Hong Kong Special Administrative Region, the Health and Medical Research Fund of Hong Kong (Project No. 08194126), and new Research Initiatives support from City University of Hong Kong (Project No. 9610497) to R.L. We thank the High Performance Computing Cluster of the City University of Hong Kong for providing computing resources. We thank Professor Zhao Xiaomin’s group at Northwest A&F University for providing wild-type TGEV and gene7-deleted TGEV.

## Notes

### Competing Interest Statement

The authors have declared no competing interest.

