## Supplemental Figures for "Nanopore direct-RNA sequencing reveals TGEV epitranscriptomic and transcriptomic landscapes modulated by gene 7"

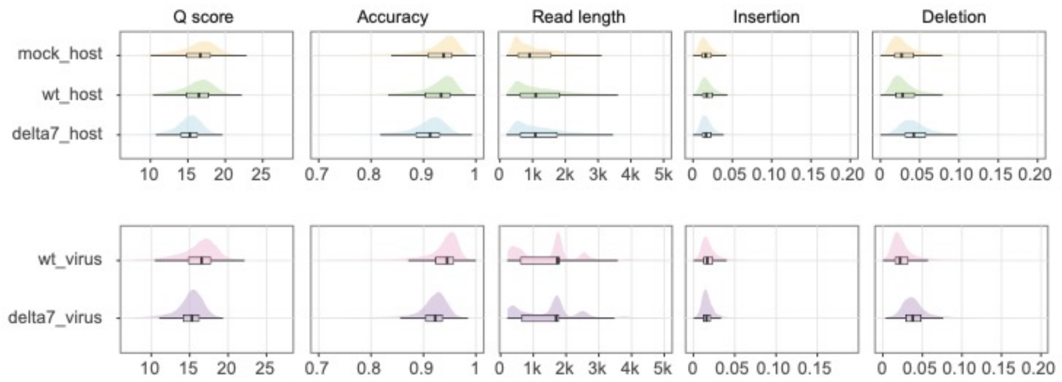


**Figure S1. Reads features of host and virus, reads basecalled using Dorado base caller.** In the upper row of figures, yellow represents mock data, green represents host data of wt, and blue represents host data of delta7. In the lower row of figures, pink represents virus data of wt while the purple represents virus data of delta7. Read-level Q score, accuracy, read length, insertions, and deletions are calculated based on the mapping results

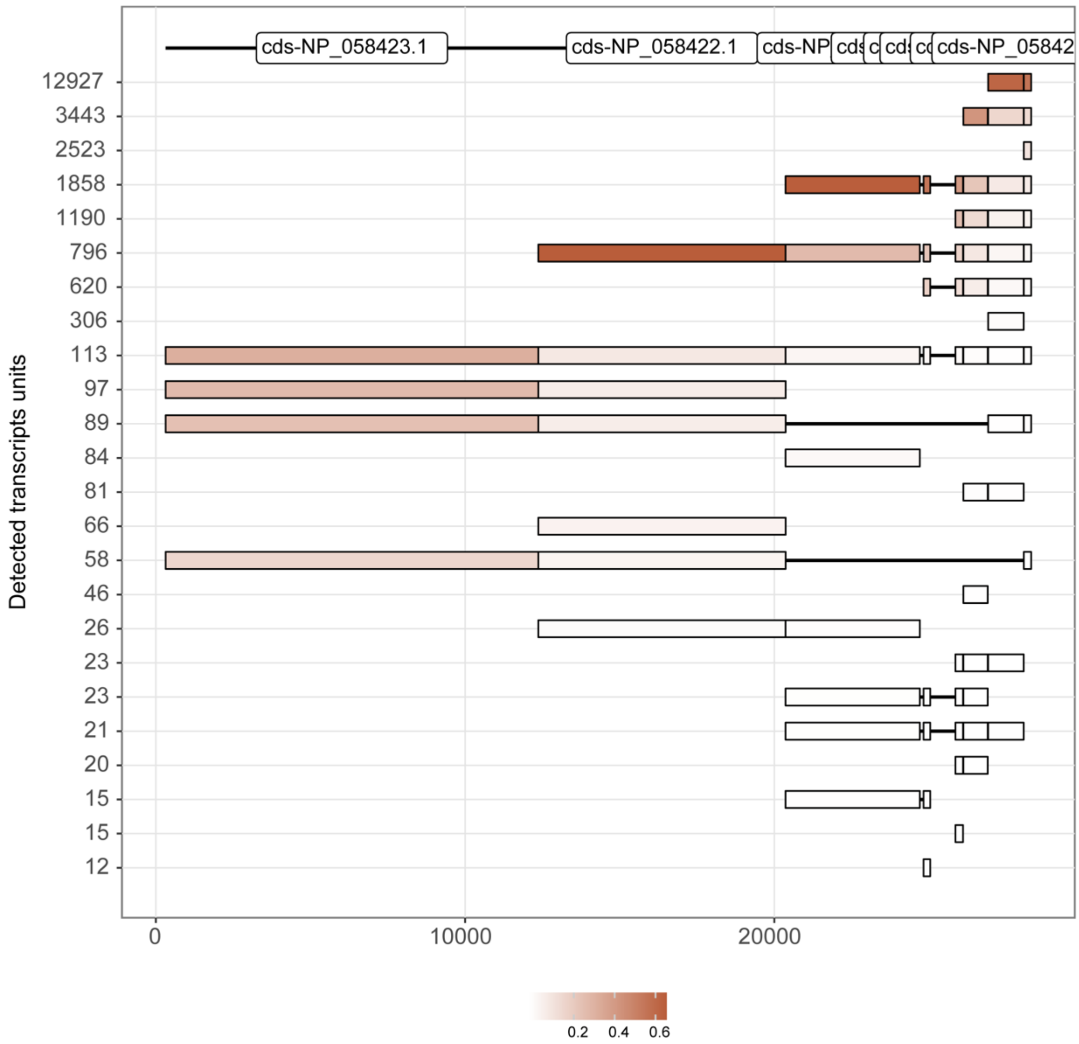


**Figure S2. Virus reads mapping result according to genomic and subgenomic RNAs.**

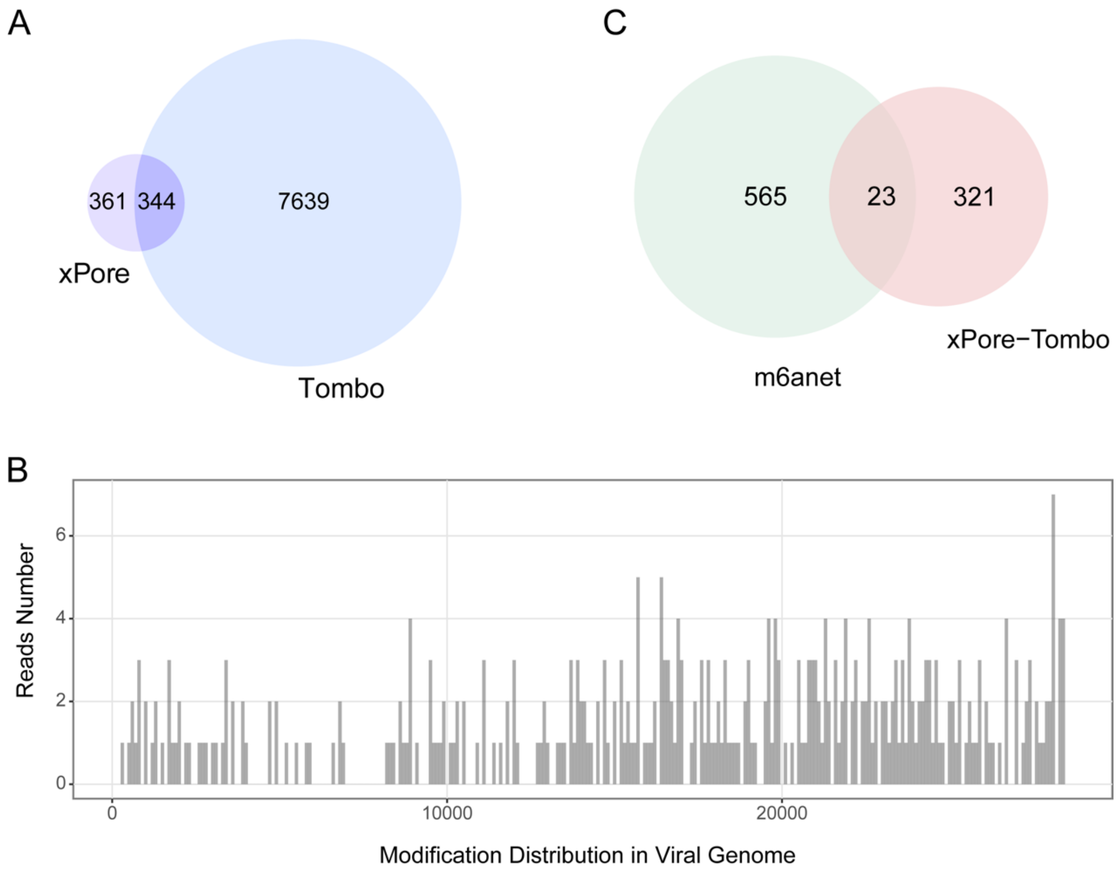


**Figure S3.** (A) Intersection of xPore and Tombo. (B) Distribution of the intersection result of xPore and Tombo. (C) Shared site of m6Anet and intersection result of xPore and Tombo.

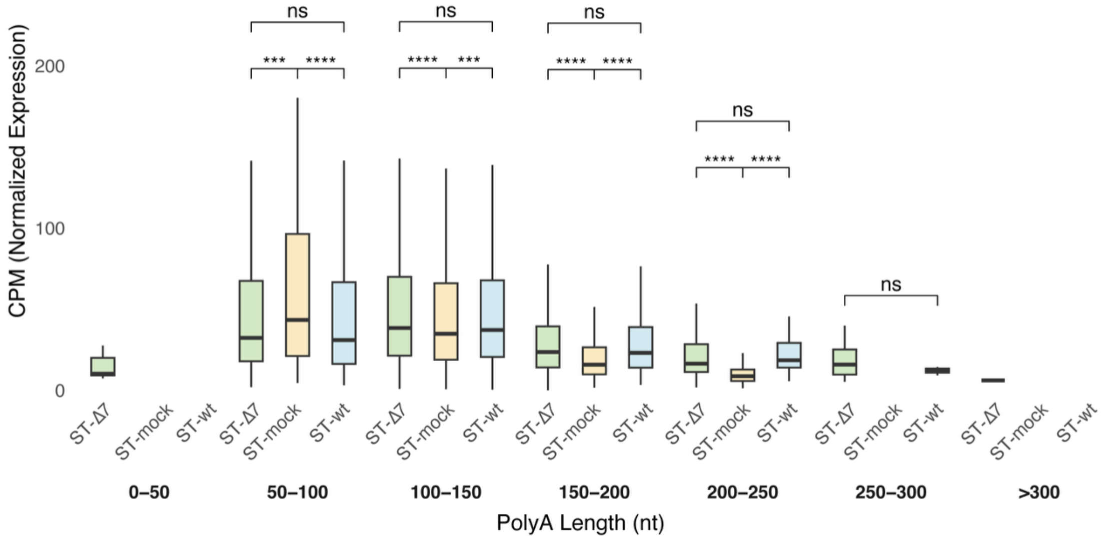


**Figure S4. Distribution of host polyA after CPM (50 nt interval)** The difference between infected hosts and healthy cells was significant. The polyA tail length of 50-100 nt was the most abundant in mock (healthy ST cells), and both ST-wt and ST-Δ7 showed more distribution in 100-150 nt. However, the loss of TGEV gene 7 did not cause a difference between ST-wt and ST-Δ7.
